# Conflict- and error-related theta activities are coupled to BOLD signals in different brain regions

**DOI:** 10.1101/2022.02.15.480552

**Authors:** Ewa Beldzik, Markus Ullsperger, Aleksandra Domagalik, Tadeusz Marek

## Abstract

Both conflict and error processing have been linked to the midfrontal theta power (4-8 Hz) increase as indicated by EEG studies and greater hemodynamic activity in the anterior midcingulate cortex (aMCC) as indicated by fMRI studies. Conveniently, the source of the midfrontal theta power was estimated in or nearby aMCC. However, previous studies using concurrent EEG and fMRI recordings in resting-state or other cognitive tasks observed only a negative relationship between theta power and BOLD signal in the brain regions typically showing task-related deactivations. In this study, we used a simultaneous EEG-fMRI technique to investigate a trial-by-trial coupling between theta power and hemodynamic activity during the performance of two conflict tasks. Independent component analysis (ICA) was applied to denoise the EEG signal and select individual midfrontal EEG components, whereas group ICA was applied to fMRI data to obtain a functional parcellation of the frontal cortex. Using a linear mixed-effect model, theta power was coupled with the peak of hemodynamic responses from various frontal, cingulate, and insular cortical sites to unravel the potential brain sources that contribute to conflict- and error-related theta variability. Although several brain regions exhibited conflict-related increases in hemodynamic activity, the conflict pre-response theta showed only a negative correlation to BOLD signal in the midline area 9 (MA9), a region exhibiting conflict-sensitive deactivation. Conversely, and more expectedly, error-related theta showed a positive relationship to activity in the aMCC. Our results provide novel evidence suggesting that the amplitude of pre-response theta reflects the process of active inhibition that suppresses the MA9 activity. This process is affected independently by the stimulus congruency, reaction times variance, and is susceptible to the time-on-task effect. Finally, it predicts the commitment of an omission error. Together, our findings highlight that conflict- and error-related theta oscillations represent fundamentally different processes.

## 1. Introduction

One of the most-studied human EEG oscillations, the midfrontal theta rhythm (4-8 Hz), has been consistently reported to increase during various cognitive tasks and was implicated to have a crucial, active role in many cognitive processes. Midfrontal theta power (MFT) increase has been linked to conflict detection and resolution (Cohen, 2014; Jiang et al., 2018), error detection and feedback processing (Cavanagh et al., 2012; Luu et al., 2003), working memory (Brzezicka et al., 2018; Onton et al., 2005), language comprehension (Bastiaansen et al., 2005) or mental arithmetic (Gärtner et al., 2015). The wide-ranging engagement of MFT in numerous cognitive tasks was inferred that theta plays a general role in cognitive control mechanisms (Cavanagh and Shackman, 2015; Duprez et al., 2020; Hanslmayr et al., 2008). Interestingly, studies employing simultaneous EEG-fMRI technique in resting-state and working memory tasks have coupled the MFT power to, and only to, cortical brain regions, which show task-related deactivations (Algermissen et al., 2021; Scheeringa et al., 2009, 2008). Would the same hold true for the MFT involved in conflict tasks, given that response conflict has been associated with increased midfrontal brain activity (Ullsperger and Von Cramon, 2001)? In the current study, we aimed to address this question.

The last two decades of neuroscience research provided significant advancement in understanding cognitive control mechanisms involved in conflict resolution and error detection. To date, response conflict tasks provide a great model to study these processes because of several behavioral effects that they elicit, e.g., the congruency effect, conflict adaptation effect (also known as the Gratton effect), conflict trial proportion effect, as well as high error likelihood (Bausenhart et al., 2021; Cespón et al., 2020; Chuderski and Smolen, 2016; Heidlmayr et al., 2020). In EEG literature, greater activity in the MFT is consistently reported for both congruency effect and error commission (Cavanagh and Frank, 2014; Nigbur et al., 2011). Conflict-related theta increase peaks in the FCz channel around 300-500 ms after the stimulus or before the response at 6 Hz frequency. Its source was estimated in the pre-Supplementary Motor Area (preSMA) using dipole fitting (Nigbur et al., 2011) or beamforming (Cohen and Ridderinkhof, 2013) tools. Its key characteristic is being non-phase-locked to stimulus onset or button press (Cohen, 2014; Cohen and Donner, 2013), yet its phase was coupled to oscillatory activity in distinct brain sites (Asanowicz et al., 2021; Cavanagh et al., 2009a; Cohen and Ridderinkhof, 2013; Hanslmayr et al., 2008). Although MFT is considered a marker of conflict processing (Nigbur et al., 2011), a recent account suggests its more general role related to response execution rather than conflict per se (Duprez et al., 2020). Conversely, error-related theta is phase-locked to the response onset, peaks in FCz channel 50-100 ms after the response at 5 Hz, and can be linked to the evoked potential called error-related negativity (Cavanagh et al., 2009b; Cavanagh and Frank, 2014; Luu et al., 2004a; Trujillo and Allen, 2007). Its source was estimated in the anterior midcingulate cortex (aMCC) and linked with action monitoring and post-error behavioral adjustments (Beldzik et al., 2015a; Debener et al., 2005; Valadez and Simons, 2018).

The fMRI experiments investigating the congruency effect were under great influence of the conflict monitoring model (Botvinick et al., 2001) due to an apparent and replicable activity increase of the aMCC for incongruent trials compared to congruent ones. However, this theory was widely debated due to a confounding factor of reaction times (RTs) effect on the aMCC activity (Carp et al., 2010; Domagalik et al., 2014; Grinband et al., 2011) and the models’ assumptions (Schmidt, 2019; Vassena et al., 2017). Regarding the neural substrate for error processing, there is little controversy. In line with the EEG source analysis, the aMCC was found a brain locus for a greater activity for erroneous trials than correct ones (Beldzik et al., 2018; Holroyd et al., 2004; Iannaccone et al., 2015; Wessel et al., 2012) in line with current aMCC accounts suggesting its central role in signaling surprise and the need for adaptive behavior (Ullsperger et al., 2014; Vassena et al., 2020). Interestingly, other fMRI studies provided evidence that the preSMA activity is sensitive to congruency effect (Iannaccone et al., 2015; Nachev et al., 2008; Ullsperger and Von Cramon, 2001) and unbiased by the RT effect (Beldzik et al., 2015b). Thus, it was indicated that the preSMA is responsible for action selection by detecting competing motor plans and inhibiting prepotent responses (Huster et al., 2011; Iannaccone et al., 2015).

The EEG and fMRI literature provide us with distinct but highly complementary information of when and where, respectively, cognitive processes occur. The best way to verify and combine the findings of electrophysiological and neuroimaging measurements is by conducting a simultaneous EEG-fMRI experiment. This technique proved already successful in coupling event-related potentials in conflict tasks with BOLD signals (Debener et al., 2005; Iannaccone et al., 2015). However, no previous study has coupled MFT activity in conflict tasks with the fMRI activity searching for the potential source of the theta generator. Considering the abovementioned literature, one would expect conflict-related theta to be generated in the preSMA, whereas error-related theta in the aMCC. However, previous studies aiming at coupling MFT with cortical fMRI activity observed only a negative relationship between these brain measures (Algermissen et al., 2021; Prestel et al., 2018; Scheeringa et al., 2009, 2008). This negative correlation was found in the midline areas 9 and 10 that are, among other parietal and temporal regions, attributed to the default mode network (Andrews-Hanna et al., 2014). These regions show task-related deactivations for most cognitive tasks, which is interpreted as supporting goal-directed behavior by suppressing task-irrelevant functions such as metacognitive thoughts, mind-wandering, or autobiographical memory retrieval (Anticevic et al., 2012; Beldzik et al., 2021; Mayer et al., 2010). Yet, the relationship between MFT and BOLD signal in conflict and error processing remains unclear.

This study aimed to fill that gap by implementing simultaneous EEG-fMRI measurements to conflict tasks. Our goal was to obtain the MFT EEG activity and couple its amplitude with the BOLD signal in the medial frontal cortex on a trial-by-trial basis. Such analysis was conducted separately for correct trials varying in congruency level (conflict analysis) and incongruent trials varying in accuracy level (error analysis). Since multiple brain regions are involved in conflict and error processing, we assumed that the activity of multiple brain regions can contribute to the MFT power variance. Despite accounting for this assumption in a linear mixed-effect model, a single theta-BOLD relationship was observed for each type of analysis. Interestingly, this relationship referred to different brain regions indicating disparate characteristics of conflict and error theta activities.

## 2. Methods

### 2.1 Participants

Thirty-seven participants (mean age, 22.1 ± 2.7 years; 22 females/15 male) met the following experiment requirements: no contraindication for MRI scanning, right-handedness (verified with the Edinburgh Handedness Inventory; Oldfield, 1971), normal or corrected-to-normal vision, no color- blindness (confirmed with Ishihara color vision test), no reported physical or psychiatric disorders, drug-free. Participants were informed about the procedure and goals of the study, and they gave written consent. The study was approved by the Bioethics Commission at Jagiellonian University. The study was carried out in accordance with the guidelines of the declaration of Helsinki and. Data from two subjects were excluded during analysis due to the lack of frontocentral components in the EEG data (see Method section 2.5).

### 2.2 Experimental task

The task was prepared and generated using E-Prime 2.0 (Psychology Software Tools). It was presented on a 32-inch screen, located behind the MR scanner and approximately 100 cm from the head coil. Participants were able to see the screen using a single mirror placed on the head coil.

Participants performed two types of conflict-inducing tasks, i.e., the Stroop (Fig. 1A) and the Simon (Fig. 1B) tasks. In the former, a stimulus was one of four color names (red, yellow, blue, or green) in Polish (Arial font, height 2°) printed in one of these colors. In the latter, a stimulus was a dot (diameter 2°) presented laterally (∼22°) to the fixation sign printed in one of these four colors. Although the tasks differed in stimulus features, they had the exact instructions given to participants: “Indicate an ink color of a stimulus ignoring its other features.” Indicating a color was obtained by pressing a specified button of response grip (Nordic Neuro Lab, Bergen, Norway) using a specified finger (left index finger, left thumb, right index finger, right thumb, respectively; Fig 1).

**Figure 1.**
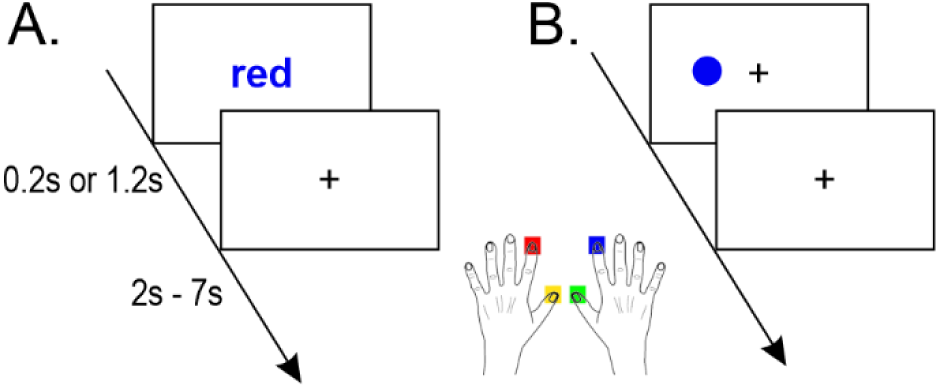
Scheme of the A) Stroop and B) Simon tasks.

Response conflict was present on trials where the target response was a semantic mismatch or contralateral to the stimulus location (“incongruent trials”). Conversely, no response conflict was present on trials where the target response was a semantic match or ipsilateral to the stimulus location (“congruent trials”). The stimulus was presented for either 200 ms (“short”) or 1200 ms (“long”). In both cases, the response window was 1.2 s. A ‘speed up’ icon was shown in case of a missing response within that time. A fixation point (a plus sign, size 2° × 2°) was present throughout the whole experiment except for the stimuli presentation period in the Stroop task (see Fig. 1). The intertrial time interval (ITI) was randomly drawn from a uniform distribution of values in the following categories: 0.8, 1.3, 1.8, 2.3, 2.8, 5.3, and 5.8 s, resulting in an average of 3.5 s.

Each task was presented in two blocks of 50% and 20% congruency rates. Each session consisted of 60 incongruent trials and 60 or 240, respectively, congruent trials. These four sessions were counterbalanced between participants, with one restriction being the task type in the interleaved fashion (e.g., Stroop20%, Simon50%, Stroop50%, Simon20%). Between the sessions, subjects had an unlimited break from the task. They were informed of the next stimulus type (word or dot) and a reminder with the response options before beginning a new session. In total, the experiment lasted approximately 50 min and introduced 840 trials. Together, our design controlled for conflict (congruent vs. incongruent), task (Stroop vs. Simon), congruency rate (50% vs. 20%), and stimulus duration (short vs. long). The rationale for including these four conditions was to account for various task parameters, which differ in EEG and fMRI protocols.

Before the main experiment, participants undertook a training paradigm, which included 12 trials of centrally presented dots (neutral trials), 12 trials of Simon, and 12 Stroop tasks. If accuracy in each 12- trial run was below 90%, a subject had to redo the run.

### 2.3 Behavioural data analysis

The ratio for erroneous responses and omissions was calculated for each participant. The former was verified in the context of the congruency trial-type using a paired t-test. Reaction times (RTs) of correct responses underwent a linear mixed-effects (LME) model, which is less prone to Type I and II errors than conventional regression methods (Aarts et al., 2014). In the model, task parameters were entered as fixed dependent variables, whereas subjects were entered as random effects. Additionally, congruency rate and type of trial were included in the model with an interaction term due to the expected intersecting pattern of these two effects (i.e., the opposite effect on RT in congruent and incongruent trials in case of high congruency rate). Thus, the LME model was captured using the following equation:

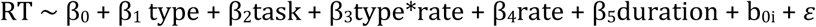

in which the RT is a function of the intercept (β_0_), trial type (β_1_), task (β_2_), the interaction of type and congruency rate (β_3_), congruency rate (β_4_), stimulus duration (β_5_), deviation of a given participant from the intercept (b_0i_), and the error term ε. The simplified version of the equation above is:

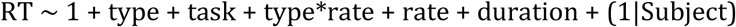

Further on, we shall use only the simplified notation.

### 2.4 EEG data acquisition and preprocessing

EEG data were recorded using an MR-compatible EEG cap (EasyCap, Herrsching, Germany) with 63 scalp electrodes, following the extended International 10-20 system and an additional channel for recording an electrocardiogram (ECG). The ECG electrode was placed on the participants’ back under the left shoulder blade to avoid signal contamination with chest movements. The reference electrode was positioned at FCz. EEG data were recorded using Vision Recorder (Version 1.20) with a sampling rate of 5 kHz. The electrode impedances were kept below 20 kΩ. The SyncBox (Brain Products GmbH, Gilching, Germany) was used to ensure that the EEG clock was synchronized to the MR scanner clock (Mullinger et al., 2008).

The first two steps of EEG data preprocessing were conducted using BrainVision Analyzer 2.0 software (Brain Products GmbH). MR artifacts were minimized using an average artifact subtraction (AAS) technique (Allen, Josephs, & Turner, 2000). Specifically, the gradient artifact was defined as a continuous interval of 1800 ms in length, beginning at the “start volume scan” marker. An artifact template was created using a sliding average of twenty-one artefactual intervals. Next, datasets were downsampled to 500 Hz. The AAS technique was also applied to correct for ballistocardiogram artifacts. Again, twenty- one pulses in the semiautomatic mode were used to create a template. Peak detection was run on the ECG channel. EEG data were then exported to EEGLAB (version 2019.1; Delorme and Makeig, 2004), excluding the ECG channel. After removing resting periods, data were bandpass filtered in a 0.5 – 35 Hz range (*eegfiltnew*) and re-referenced by common average.

### 2.5 EEG component selection

The EEG data were temporarily epoched to mark ‘bad’ epochs. The criteria for a ‘bad’ epoch included either greater than 3 absolute normalized channel-mean variance in the period -500 ms to 1200 ms after the stimulus onset (Nolan et al., 2010) or greater than 150 μV absolute amplitude in ± 100 ms relative to the response onset, accounting for possible movement-related artifacts (Beldzik et al., 2019). These criteria marked 8.0% (SD 8.1%) of the trials.

We used the ICA denoising approach to obtain a clearly detectable frontocentral component for each participant (Beldzik et al., 2019; Scheeringa et al., 2016, 2011). First, continuous EEG data were bandpass filtered in the 4-8 Hz frequency range. Following epoch extraction in 0-1200 ms poststimulus range as well as the exclusion of ‘bad’ epochs and those with incorrect responses, the ICA was performed using the default, extended infomax algorithm (Lee et al., 1999). The unmixing weights were applied to the preprocessed data (i.e., before the denoising), and components were back-projected to the channel level. Applying the weights back on the original data enables performing final time-frequency decomposition on full power spectrum and extended epochs instead of theta-filtered short ones. The topographical maps were visually inspected for the most prominent frontocentral component. Out of 37 participants, two did not show any frontocentral components; thus, these subjects were removed from further analyses. Epochs were extracted from −1 s to 1.8 s relative to the stimulus presentation from the selected components’ time courses.

### 2.6 EEG time-frequency analysis

First, we aim to verify conflict-related theta increase conventionally. The time-frequency decomposition was carried out using the complex Morlet wavelet convolution. The frequency vector comprised 60 points in the 2-30 Hz range, increasing logarithmically. Similarly, the cycle values corresponding to each frequency were in a range of 2-7, increasing logarithmically. The calculated spectral power was baseline-corrected by subtracting the mean power -500 to -200 ms before stimulus onset from each time point (Duprez et al., 2020). Next, the time-frequency plots were epoched from 0 to 1200 ms aligned to the stimulus onset (stimulus-locked data) and from -800 to 400ms aligned to the response onset (response-locked data) in 20ms resolution. The maps were averaged and compared using a paired two- tailed t-test separately for conflict (correct congruent vs. correct incongruent trials) and error (erroneous incongruent vs. correct incongruent trials) analysis types. The *p* values corresponding to each pixel of the differential maps were corrected with a false discovery rate (FDR) at α < .01.

### 2.7 fMRI data acquisition

MRI was performed using a 3T scanner (Magnetom Skyra, Siemens) with a 64-channel head/neck coil. The isocenter was set 4 cm superior to the nasion to reduce gradient artifacts in the EEG data (Mullinger et al., 2011). High-resolution, whole-brain anatomical images were acquired using a T1-MPRAGE sequence. A total of 176 sagittal slices were obtained (voxel size 1 × 1 × 1.1 mm3; TR = 2,300 ms, TE = 2.98 ms, flip angle = 9°) for coregistration with the fMRI data. Next, a B0 inhomogeneity gradient field map (magnitude and phase images) was acquired with a dual-echo gradient-echo sequence, matched spatially with fMRI scans (TE1 = 4.92 ms, TE2 = 7.38 ms, TR = 508 ms).

Functional T2*-weighted images were acquired using a whole-brain echo-planar (EPI) pulse sequence with the following parameters: 3.5 mm isotropic voxel, TR = 1800ms, TE = 27 ms, flip angle = 75°, FOV 224 × 224 mm^2^, GRAPPA acceleration factor 2, and phase encoding A/P. Whole-brain images (cerebellum excluded) were covered with 34 axial slices taken in an interleaved order. Due to magnetic saturation effects, each session’s first three volumes (dummy scans) were discarded instantly, resulting in two sessions of 240 and two sessions of 590 volumes acquired for each participant.

### 2.8 fMRI data preprocessing

Preprocessing of fMRI data was conducted using Analysis of Functional NeuroImage (AFNI, version 17.3.03; Cox, 1996) and the FMRIB Software Library (FSL, version 5.0.9; Jenkinson et al., 2012). Anatomical images were skull-stripped and coregistered to MNI (Montreal Neurological Institute) space using nonlinear transformation (*@SSwarper*). They were segmented (*FAST*) to create individual cerebrospinal fluid (CSF) masks. The first step of functional data preprocessing was to obtain the transformation matrix for motion correction (*3dvolreg*) to avoid its possible alteration by temporal interpolation applied further to fMRI data (Power et al., 2017). Next, de-spiking (*3dDespike*) and slice timing correction (*3dTshift*) were conducted. Then, transformation matrices for coregistration of functional data to anatomical data (*align_epi_anat*.*py*) as well as B0 inhomogeneity derived from gradient fieldmaps (*Fugue*) were calculated. The spatial transformation was performed in one step (*3dNwarpApply*), combining all prepared matrices, i.e., motion correction, anatomical co-registration, and distortion correction. The fMRI datasets were masked using a clip level fraction of 0.4, scaled to percent signal change, and the CSF signal was extracted using previously obtained individual masks. Finally, the functional images were coregistered to MNI space using the transformation matrix from nonlinear anatomical normalization.

To clear the fMRI signal from motion residuals, we applied ‘null’ regression (*3dREMLfit)* with the pre-whitening option (using ARMA_(1,1)_ model) to functional images. The model included twelve movement parameters (six demeaned originals and six first derivatives), the CSF time course, and four or nine polynomials as determined automatically using 1 + int(D/150) equation, where D is the session’s duration. The rationale for regressing out the CSF signal lies in the fact that this signal reflects purely physiological noises, respiratory and cardiac, and often contains motion-related artifacts (Caballero-Gaudes and Reynolds, 2017; Power et al., 2014).

### 2.9 fMRI data analysis

Our interim goal was to obtain a ‘functional parcellation’ of the BOLD signal in the frontal cortex to verify all potential MFT source(s) precisely. Thus, a group ICA was conducted (GIFT version 4.0b; (Calhoun et al., 2001)Calhoun et al., 2001) using a mask limited to all frontal, insular, and cingulate regions defined by the Harvard-Oxford cortical structural atlas (neurovault id: 1705). An estimation to determine the number of components was performed using minimum description length (MDL) criteria (Y.-O. Li et al., 2007). ICA decomposition stability was validated using ICASSO(Himberg et al., 2004) with 50 random initializations of the Infomax algorithm. Data were back-reconstructed using the default GICA option with z-scoring applied to both maps and time-courses. The components’ maps were corrected with FDR at α < .01 and inspected to identify and discard those primarily associated with artifacts representing signals from large vessels, ventricles, motion, and susceptibility (Griffanti et al., 2017; Kelly et al., 2010; Varoquaux et al., 2010).

The time course of the brains’ components was interpolated to 100-ms resolution. Next, epochs were extracted from 0 s to 10 s of the stimulus onset and baseline corrected by subtracting the values at 0. For each of the brains’ components, these fMRI epochs were averaged for each and compared using a paired two-tailed t-test separately for conflict (correct congruent vs. correct incongruent trials) and error (erroneous incongruent vs. correct incongruent trials) analysis types. The *p* values corresponding to each time-point of the hemodynamic responses were corrected with FDR at the α < .01.

### 2.10. Integration of EEG-fMRI data

Similar to RT analysis, EEG and fMRI data integration was carried out using LME. The model is ideal for coupling two experimental measures as it allows including single-trial measurements from all subjects in one group analysis (Beldzik et al., 2019; Wichary et al., 2017). In the model, theta values were entered as a dependent variable, whereas the hemodynamic BOLD activity was entered as a predictor. Although the hemodynamic response has a major delay, it reflects brain activity that constitutes a potential source of scalp-registered theta amplitudes. Moreover, there may be multiple sources of frontocentral theta, which can be verified by entering multiple fMRI components into the model. Furthermore, an accurate trial selection can modulate predictor sensitivity to experimental conditions. Here, all correct trials and all incongruent trials were selected to account for conflict-related and error-related variance, respectively. Finally, the model controls other factors affecting both neural measures, such as ITI and block number. Thus, LME comprises a comprehensive and flexible tool to explore a potential source of the theta activity. Such analysis was conducted in two ways - a single and multiple fashions, as explained below.

The first approach was to apply a single LME model to theta values extracted from the window of interest separately for correct trials varying in congruency level (conflict analysis) and incongruent trials varying in accuracy level (error analysis). In the conflict analysis, the mean amplitude in the theta range (4-8 Hz) was calculated for all correct trials within the stimulus epoch (0-1200 ms in the stimulus- locked data) and within the conflict-sensitive time window (determined based on the time-frequency EEG results). Additionally, maximum values in the theta band were extracted from the stimulus epoch window. The rationale for these various extraction protocols was the possibility that different brain regions contribute to different aspects of theta activity, e.g., tonic (epoch mean value) or phasic (maximum value in the epoch). In the error analysis, the mean amplitude in theta range was calculated for all incongruent trials within the post-response epoch due to theta’s high sensitivity to accuracy score in that period. Similar to conflict analysis, the length of the theta extraction window was determined based on the time-frequency EEG results for the accuracy condition.

The BOLD values at the peak of hemodynamic activity, i.e., mean activity between 3-5 s after stimulus onset, were extracted for each brain component. Due to the high statistical power of the single model, it was possible to enter all brain components as predictors. The rationale behind this decision was the possibility that multiple cortical sources contribute to the midfrontal EEG signal due to volume conduction. Furthermore, the model accounts for possible residual autocorrelation between fMRI components, distinguishing those that explain most of the theta variance without running multiple tests. Additionally, the whole-epoch mean power in the remaining EEG frequency bands, i.e., delta (2-3 Hz), alpha (9-12 Hz), and beta (13-30 Hz), was implemented as fixed effects to control for the spectral leakage and the possible dominant activity in the other frequency bands. Subjects’ number, ITI category, and block number were entered as random effects in this model. This way, assuming random intercepts allowed us to account for each category’s deviation from the average intercept in the model.

The final matrix comprised 24326 and 7291 observations in conflict and accuracy analyses, respectively. All continuous observations were z-scored for each subject before running the test. The simplified notion of the *single model* was as follows:

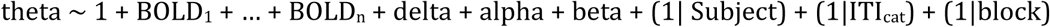

Additionally, we verified if these extracted theta values remain sensitive to conflict or accuracy analyses while controlling for RT variability. Thus, another LME model was implemented, in which theta was a function of trial type and individually z-scored RT values. For coherency reasons, the same random effects were used.

The second approach was to apply the LME model to each frequency power at each time point. Such analysis aims to confirm the single model results and determine the precise timing of the relationship. These multiple models were run similarly to the single one. However, due to lower statistical power, a BOLD signal from only one fMRI component was entered into the equation. Therefore, LME was run n- times for each brain region, resulting in the time-frequency maps of parameter estimates between oscillatory EEG power and BOLD signal. The simplified notion of the *multiple models* was as follows:

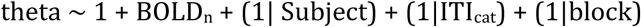

Only response-locked time-frequency maps underwent these computations. To save computational time, the maps were downsampled to 30 frequency bins (2-30 Hz range) and 50 ms time resolution (from -400 ms to 300 ms). The maps were corrected with a false discovery rate (FDR) at the α<.01.

### 2.11 Follow-up analysis

Additionally, two follow-up analyses were conducted to better characterize the brain measures obtained here and strengthen the inferences about their role in the task. First, the amplitude of EEG spectral power and fMRI hemodynamic responses underwent a traditional subject-level comparison between omission and correct trials. These analyses were conducted similar to conflict and accuracy comparisons except that no baseline was provided here due to expected effects before or at the early stage of stimulus processing. Additionally, EEG spectral power was calculated to decibels, mean values in theta/alpha range in the 1s-long prestimulus time window were extracted and compared using a paired two-tailed t-test. The length of the window and the frequency range were determined based on the obtained time-frequency EEG maps.

Second, the epoch-mean theta activity and the peak of brain fMRI components underwent LME models to account for the time-on-task effect. Particularly,

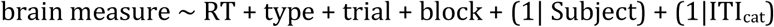

where *trial* represents the number of trials within each block session and *block* represents the number of block sessions. All variables in the model were z-scored for each subject. Additionally, a similar model was applied to RTs, which were predicted with the type, trial, and block fixed effects.

## 3. Results

### 3.1 Behavioral results

Participants (N=35) committed 5.9% (SD 3.9%) erroneous responses and 3.9% (SD 3.2%) omissions. Erroneous responses were committed twice as often on incongruent trials as on congruent trials (con: 4.7%; incon: 8.8%; t_(34)_ = 7.7; p < .001). Further behavioral analyses were conducted on correct trials only. The RTs underwent LME with all experimental conditions as fixed effects, i.e., conflict, task, congruency rate, stimulus duration (see Methods for details), and subjects as a random effect. The mean reaction time was 674.0 ms and was significantly affected by all conditions (Table 1; Fig 2). Reactions were slower by 68.5 ms in the case of incongruent trials in comparison to congruent ones. They were slower by 43.1 ms in the Stroop task in comparison to the Simon task. Responses to incongruent trials in blocks with a 20% congruency rate were slower by 32.2 ms in comparison to congruent trials in the 50% rate sessions. Finally, a minor increase of RTs was observed for 50% congruency rate and short stimulus duration (i.e., 200 ms). All these effects were in accordance with the literature findings.

**Table 1.** Results of the LME model applied to RT variance.

| Effect | Estim | SE | t | DF | pValue | Lower | Upper |
| --- | --- | --- | --- | --- | --- | --- | --- |
| Intercept | 674.0 | 9.8 | 68.6 | 24394 | 0.001 | 654.7 | 693.2 |
| type_incon | 68.5 | 3.3 | 20.9 | 24394 | 0.001 | 62.1 | 74.9 |
| task_Stroop | 43.1 | 1.8 | 24.6 | 24394 | 0.001 | 46.5 | 39.6 |
| type_incon:rate_20% | 35.1 | 4.2 | 8.3 | 24394 | 0.001 | 26.8 | 43.4 |
| rate_20% | -12.0 | 2.6 | -4.7 | 24394 | 0.001 | -17.0 | -7.0 |
| dur_long | -4.5 | 1.8 | -2.5 | 24394 | 0.011 | -7.9 | -1.0 |

Note. Estim = parameter estimate. SE = standard error. DF = degrees of freedom. Lower/Upper = lower/upper confidence interval.

**Figure 2.**
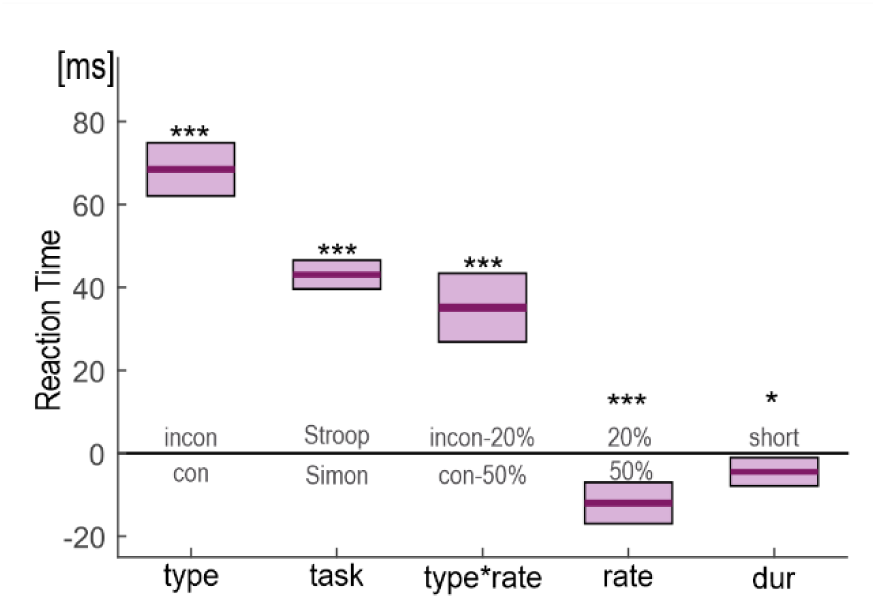
Behavioral results. Parameter estimates of LME model explaining RT variance. Incon = incongruent trials. con = congruent trials. *** p <.001; * p <. 05.

### 3.2 EEG results

The selected EEG components with frontocentral topography showed a maximum at the FCz channel (Fig. 3A). Time-frequency analyses applied to their time-courses resulted in an expected pattern of increased theta activity in high incongruency condition (Cohen and Donner, 2013; Nigbur et al., 2011). Correct incongruent trials evoked greater amplitude power in the theta band than congruent trials (Fig. 3B). This difference was most pronounced in the response-locked window from -300 ms before the response until the response itself. Erroneous trials evoked greater theta power in the 300 ms window beginning immediately after the response. In accordance with the previous studies, the peak of conflict theta was 6.3 Hz, whereas the peak of error-related theta was 4.6 Hz. Thus, the EEG results obtained here confirmed the occurrence of conflict- and error-related theta activities as both the timing and peak frequency of these effects are similar to those previously reported (Nigbur et al., 2011; Trujillo and Allen, 2007). Based on these results, the time windows for conflict- and error-related theta extraction were defined as 300 ms before the response (conflict analysis) and 300 ms after the response (error analysis), respectively.

**Figure 3.**
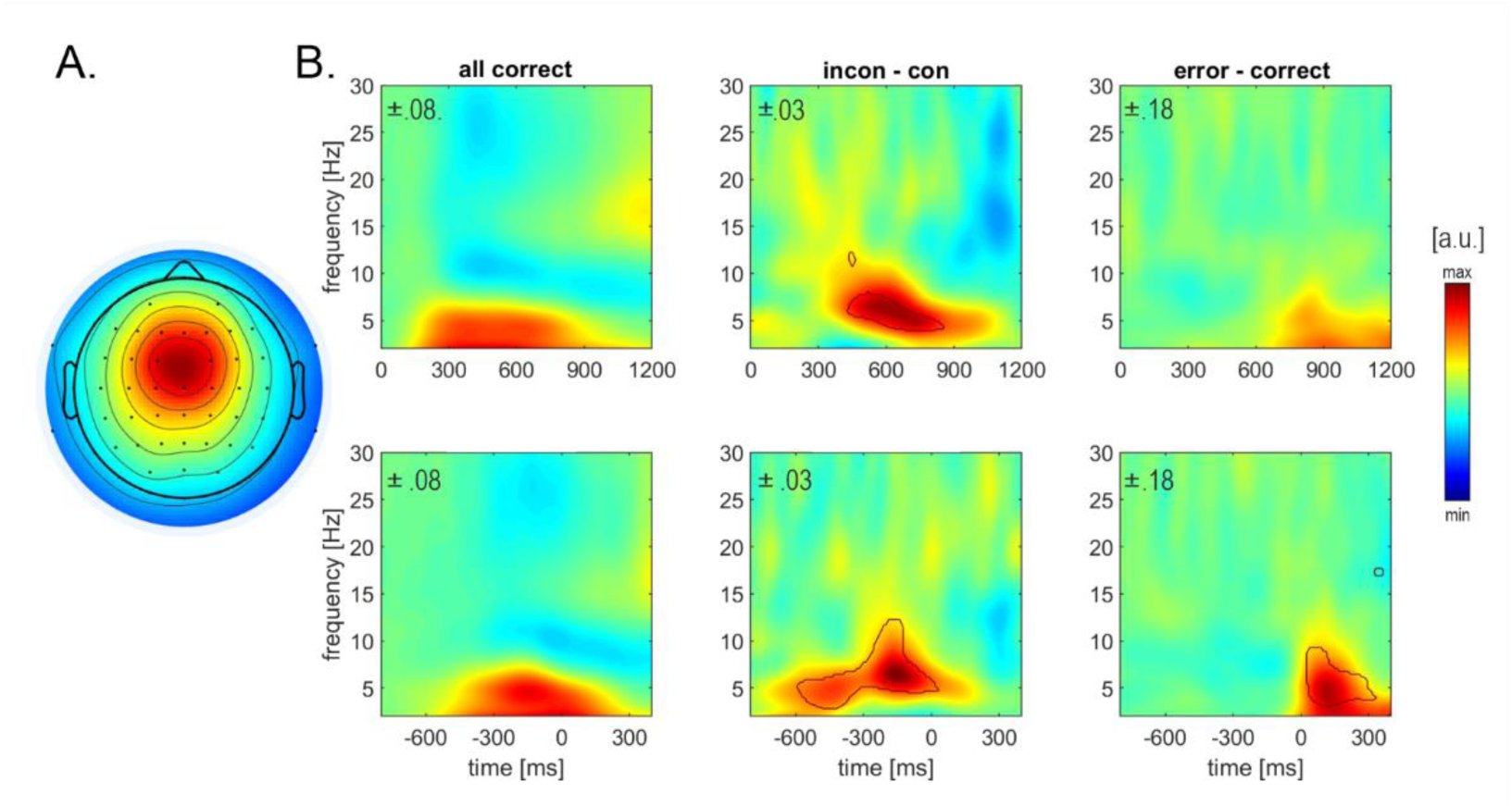
EEG results. A) Subject-mean topography of frontocentral EEG component. B) Time-frequency maps of amplitude power in stimulus-locked data (upper row) and response-locked data (lower row) for all trials, conflict, and error analyses. Black contour marks clusters of significant difference (pcor < 0.01).

### 3.3 fMRI results

The interim goal of the study was to obtain a functional parcellation of the medial frontal cortex to account for any possible source of theta activity. Although MDL criteria estimated 20 independent components, this number was raised to 25 to obtain maximal separation of the brain regions. Thus, the decomposition of ICA was conducted for 25 components using a mask covering only frontal, insular, and cingulate cortices (Fig. 4A). Based on the components’ topographies, eight of them were classified as brain regions or networks (Table 2; Fig. 4B). Their mean stability was estimated at 0.84, ranging from 0.73 to 0.94. For the labeling, we used nomenclature introduced by Sallet and colleagues (Sallet et al., 2013), focusing on their coordinates in the medial frontal cortex. Six of them, including SMA, preSMA, aMCC, midline area 8 (MA8), anterior insular cortex (AIC), and posterior insular cortex (PIC), were activated in the task, whereas the remaining two, i.e., midline area 9 (MA9) and 10 (MA10), were deactivated. The latter can be considered one of the core regions of the DMN. SMA, a part of the motor network, presented strong BOLD activity regardless of the trial type and accuracy score. PreSMA, aMCC, area 8, and AIC showed greater activity for incongruent trials and far greater for erroneous incongruent trials in comparison to congruent ones (marked with light grey shading in Fig. 4B, p_cor_ < 0.01). MA9 and MA10 showed greater deactivation only for the conflict condition.

**Table 2.** Brain regions corresponding to fMRI brain components.

| Label | Region | Side | x | y | z | t |
| --- | --- | --- | --- | --- | --- | --- |
| SMA | supplementary motor area | M | 0 | -4 | 57 | 16.1 |
|  | postcentral gyrus | R | 42 | -21 | 54 | 17.2 |
|  |  | L | -28 | -39 | 64 | 11.6 |
| preSMA | pre-supplementary motor area | M | -4 | 24 | 50 | 15.7 |
|  | inferior frontal sulcus | L | -49 | -10 | 33 | 16.1 |
|  |  | R | 49 | 31 | 22 | 13.3 |
| aMCC | cingulate gyrus | M | 3 | 24 | 33 | 16.7 |
| MA8 | medial frontal gyrus | M | 3 | 35 | 43 | 15.7 |
|  |  | R | 38 | 56 | 5 | 14.9 |
| MA9 | medial frontal gyrus | M | -3 | 45 | 40 | 14.9 |
|  | inferior frontal gyrus | L | -56 | 24 | 8 | 16.6 |
| MA10 | medial frontal gyrus | M | 0 | 59 | -2 | 12.6 |
| AIC | anterior insular cortex | L | -45 | 15 | -2 | 16.2 |
|  |  | R | 39 | 24 | -8 | 14.4 |
| PIC | insular cortex | R | 42 | -4 | 5 | 17.8 |
|  |  | L | -42 | -18 | 12 | 15.9 |

Note. R=right; L=left; M=medial; A = area; x.y.z coordinates are provided in MNI space. SMA=supplementary motor area. preSMA=pre-supplementary motor area. aMCC=anterior midcingulate cortex. AIC=anterior insular cortex. PIC=posterior insular cortex.

**Figure 4.**
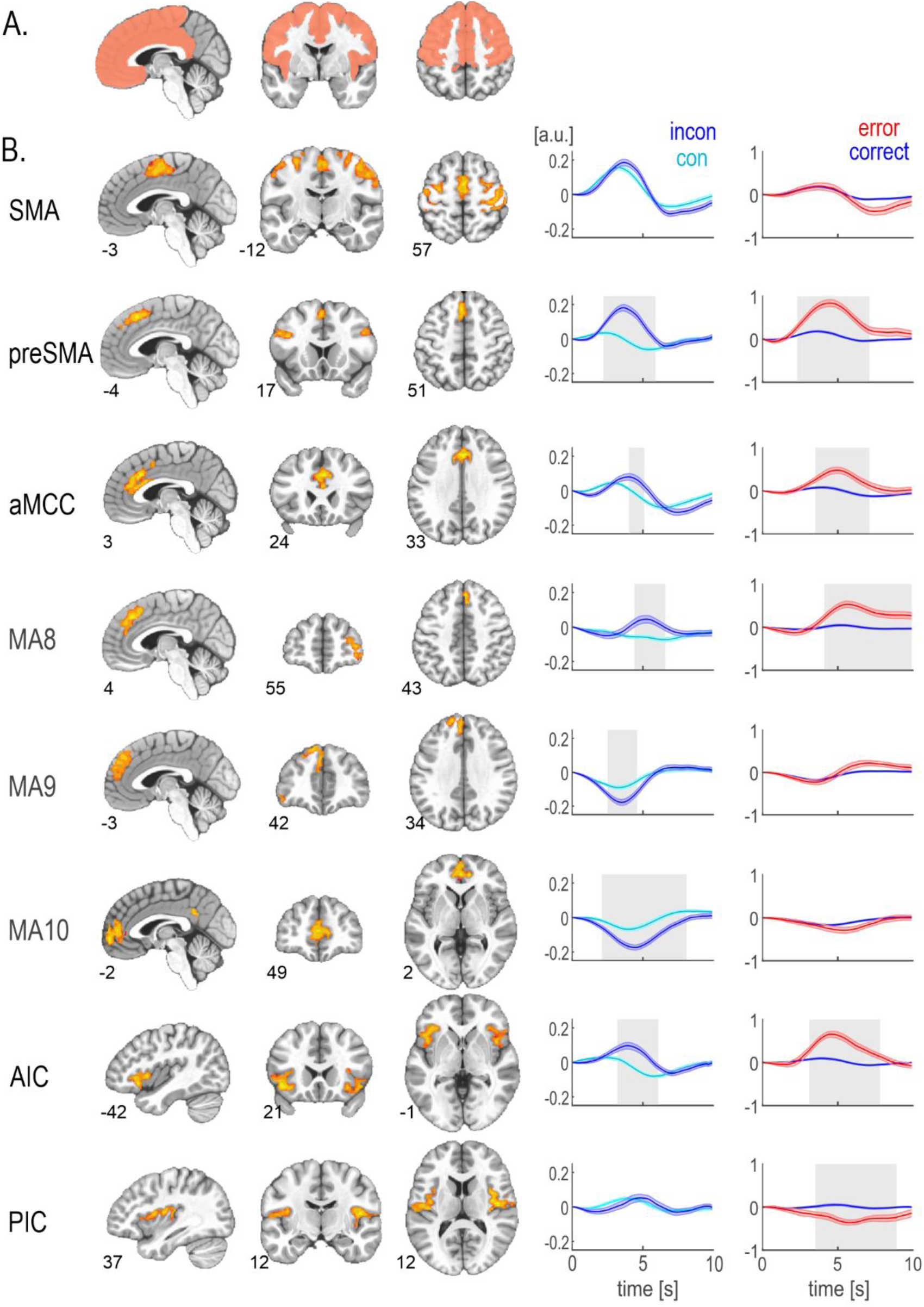
fMRI results. A) Mask covering frontal and insular cortices used in the group ICA analysis.B) Maps and averaged time-courses of brain regions or networks. The shaded area around the lines represents standard errors. A light grey area indicates a significant difference between the conditions (p_cor_ < 0.01). SMA=supplementary motor area. preSMA=pre-supplementary motor area. aMCC=anterior midcingulate cortex. AIC=anterior insular cortex. PIC=posterior insular cortex.

### 3.4 EEG-fMRI results

In the single-model approach, EEG and fMRI data integration was conducted for *a priori* pre-defined window-of-interest. In the conflict analysis, all correct trials were selected, and various time frames were used. Theta power was extracted for the whole-epoch mean, the maximum value in the epoch, and 300 ms pre-response windows. In each case, theta values proved to be significantly affected by both the congruency condition, being greater for incongruent than congruent trials, and RT variance, being greater for more prolonged reactions (Table 3; Fig 5A). The maximal theta values showed the strongest dependence on RT variance, whereas pre-response theta was the most sensitive to the conflict condition. Results of the LME model applied to fMRI data were consistent regardless of the theta extraction method (Table 3; Fig 5B). In each case, only the parameter estimates of MA9 showed significant negative values indicating a reverse relationship between BOLD amplitude and theta power.

**Table 3.** Results of a single-model approach.

| Analysis | Theta | Predictor | Estim | SE | t | DF | pValue | Lower | Upper |
| --- | --- | --- | --- | --- | --- | --- | --- | --- | --- |
| conflict | mean epoch | type_incon | 0.054 | 0.001 | 3.6 | 24323 | .001 | 0.025 | 0.084 |
|  |  | norm RT | 0.055 | 0.001 | 8.2 | 24323 | .001 | 0.042 | 0.068 |
|  |  | MA9 | -0.017 | 0.007 | -2.4 | 24314 | .016 | -0.032 | -0.003 |
|  | max epoch | type_incon | 0.043 | 0.001 | 2.9 | 24323 | .004 | 0.014 | 0.072 |
|  |  | norm RT | 0.086 | 0.001 | 12.9 | 24323 | .001 | 0.073 | 0.099 |
|  |  | MA9 | -0.028 | 0.008 | -3.7 | 24314 | .001 | -0.043 | -0.014 |
|  | pre-resp | type_incon | 0.066 | 0.001 | 4.5 | 24323 | .001 | 0.037 | 0.096 |
|  |  | norm RT | 0.052 | 0.001 | 7.8 | 24323 | .001 | 0.039 | 0.065 |
|  |  | MA9 | -0.022 | 0.008 | -2.9 | 24314 | .004 | -0.037 | -0.007 |
| error | post-resp | accur_error | 0.562 | 0.040 | 14.0 | 7288 | .001 | 0.484 | 0.641 |
|  |  | aMCC | 0.039 | 0.013 | 3.1 | 7279 | .002 | 0.014 | 0.064 |

Note. Behavioral and brain measures were implemented in separate models. Only significant estimates are presented. Estim = parameter estimate. SE = standard error. DF = degrees of freedom. Lower/Upper = lower/upper confidence interval.

**Figure 5.**
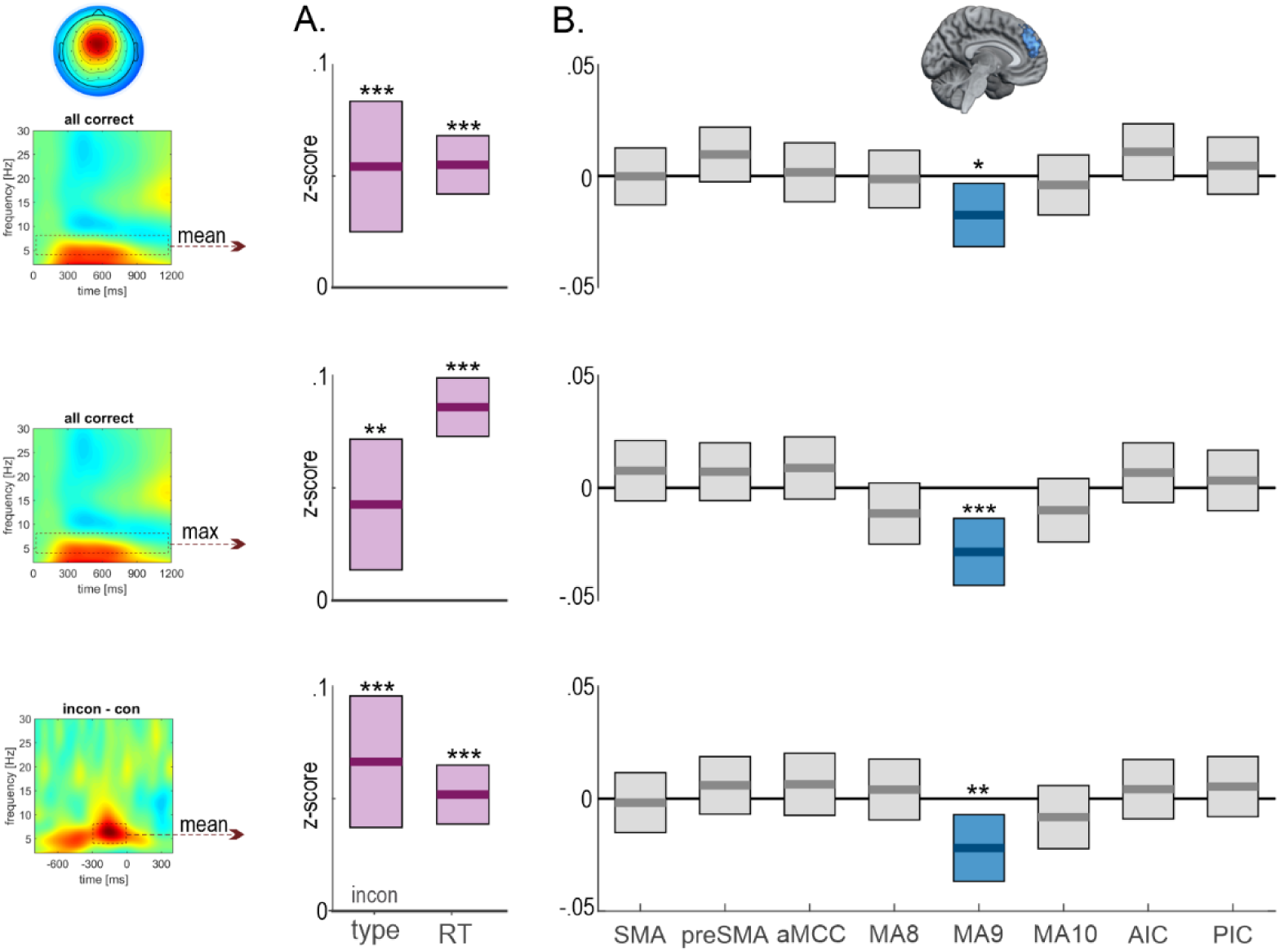
Results of a single-model approach applied to all correct trials (conflict analysis). Theta was extracted from various time windows marked with a dashed line on the time-frequency plots. Parameter estimates of theta dependency on A) type of trial, RT, and B) peak of BOLD hemodynamic responses. Horizontal bars indicate estimate values, whereas boxplots present confidence intervals. Incon = incongruent trials. Con = congruent trials. SMA=supplementary motor area. preSMA=pre-supplementary motor area. aMCC=anterior midcingulate cortex. MA=midline area. AIC=anterior insular cortex. PIC=posterior insular cortex. *** p <.001; ** p <.005; * p <.05.

In the accuracy analysis, all incongruent trials were selected. Theta power was extracted in the 300 ms-long window beginning after the response. The LME model confirmed strong dependency of these post-response theta values on accuracy score and lack of RT dependence (Table 3; Fig 6A). The LME model applied to fMRI data showed a significant positive relationship of theta with hemodynamic responses in aMCC (Table 3; Fig 6B).

**Figure 6.**
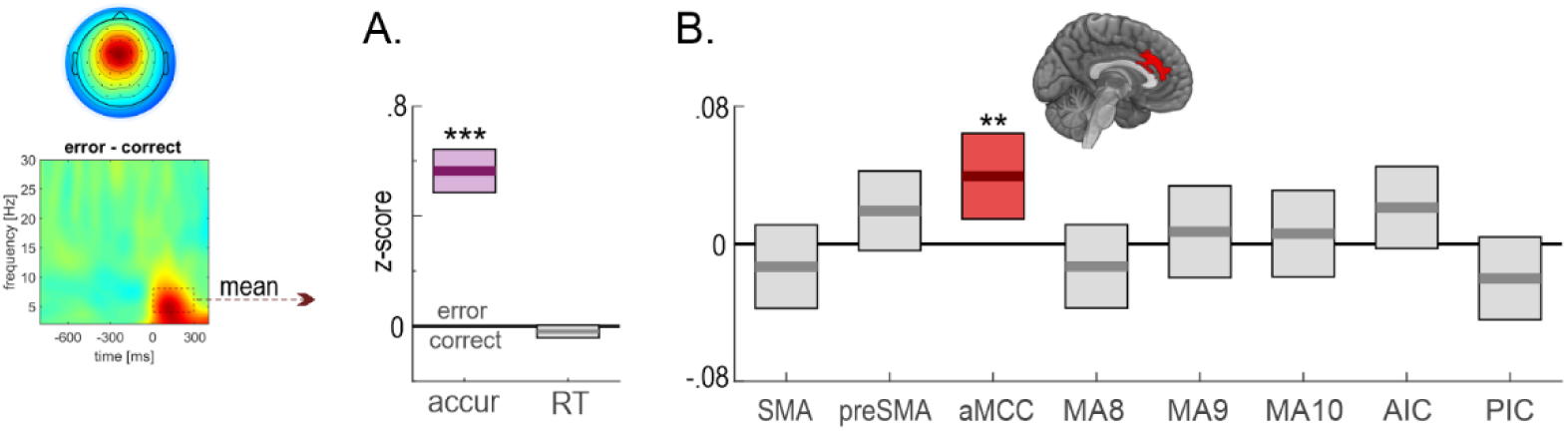
Results of a single-model approach applied to only incongruent trials (accuracy analysis). Theta was extracted from a post-response window. Parameter estimates of theta dependency on A) accuracy score, RT, and B) peak of BOLD hemodynamic responses. Horizontal bars indicate estimate values, whereas boxplots present confidence intervals. accur=accuracy. SMA=supplementary motor area. preSMA=pre-supplementary motor area. aMCC=anterior midcingulate cortex. AIC=anterior insular cortex. PIC=posterior insular cortex. *** p <.001; ** p <.005.

In the multiple-model approach, EEG and fMRI data integration was verified for each frequency bin in each time point of the time-frequency plots. Using a simplified LME model (with only one brain region at a time), we calculated the dependency between the oscillatory power and peak activity in that region separately for each analysis type. Due to substantially lower statistical power in the error analysis (three times fewer observations than in the conflict analysis), the threshold for FDR correction was reduced to α<.05.

As a result, one brain region showed a significant relationship between EEG power and BOLD signal for each analysis. Both findings were consistent with the results of the single-model approach. In the conflict analysis, a negative correlation of EEG power and hemodynamic activity was found for the MA9 (p_cor_<0.01; Fig. 7A). The most negative estimate value was observed for 5.1 Hz at 275 ms before the response occurred. In the accuracy analysis, a positive correlation of EEG power and the BOLD signal was found for the aMCC region (p_cor_<0.05; Fig. 7B). The highest estimate value was observed for 6.7 Hz at 100 ms after the response.

**Figure 7.**
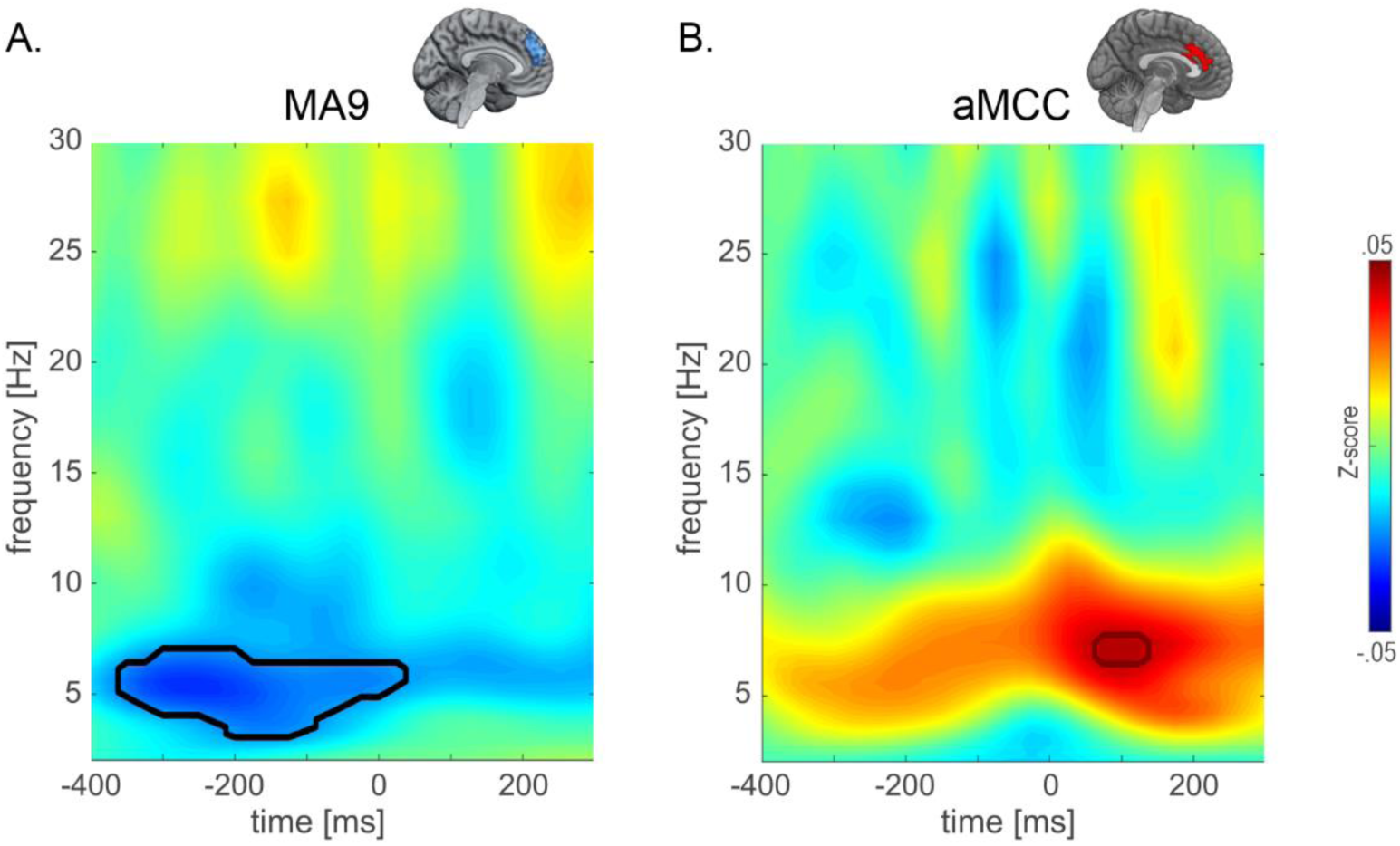
Results of a multiple-model approach applied to A) midline area 9 (MA9) activity for all correct trials (conflict analysis) and B) anterior midcingulate cortex (aMCC) activity for only incongruent trials (error analysis). Each pixel on the map presents a parameter estimate value representing the relationship between oscillatory power and peak hemodynamic response activity. Black contour marks clusters with significant values (pcor < 0.05).

### 3.5 Results of the follow-up analyses

The negative relationship between conflict theta and MA9 prompted us to conduct follow-up analyses inspired by previous findings regarding these brain measures. First, the MA9, among other default mode regions, was shown to activate preceding trials with omission error - a phenomenon linked to mind wandering (Durantin et al., 2015; Li et al., 2007). In this study, subjects committed 3.9% omissions in the task, which enabled us to replicate this observation and apply it to EEG data. The results revealed a significant prestimulus decrease (t_34_=2.04, p<0.05) of amplitude in the theta/alpha range for omissions compared to correct trials (Fig. 8A, left panel). In line with Durantin and colleagues, MA9 showed a significant increase around the stimulus onset for the same comparison (Fig. 8A, right panel).

**Figure 8.**
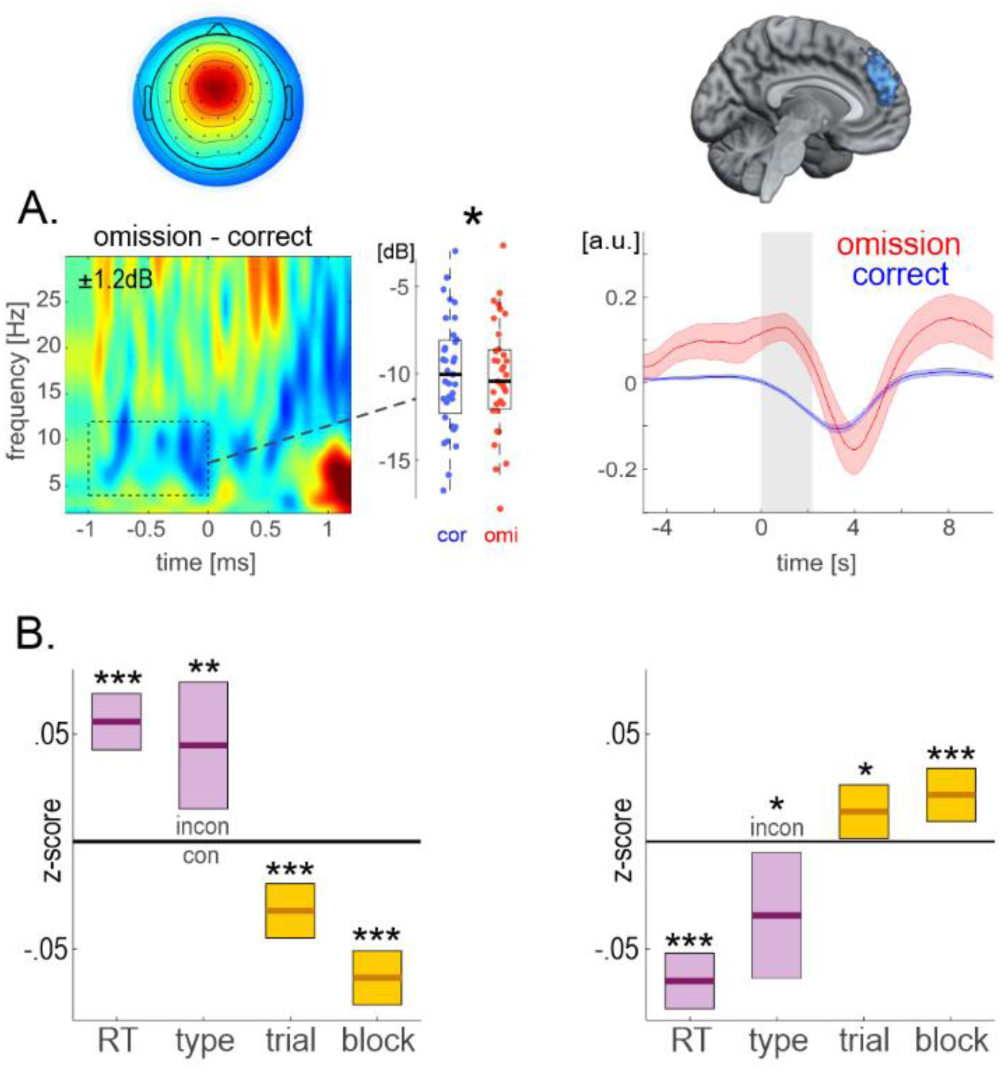
Results of the follow-up analyses. A) Comparison of the omission and correct trials for the subject-mean stimulus-locked spectral power of EEG frontocentral component (left) and hemodynamic responses in MA9 (right). Boxplot represents mean and st. dev. Cor = correct. Omi = omissions. B) Estimates for the LME model predicting mean-epoch theta values (left) and mean MA9 values around its trough (right) as a function of major behavioral and time-on-task effects. Horizontal bars indicate estimate values, whereas boxplots present confidence intervals. RT = reaction times. Incon = incongruent trials. Con = congruent trials. *** p <.001; ** p <.005; * p <.05.

Second, a recent study by Arnau and colleagues (2021) pointed to a sustained decline in task-related MFT activity with the time spent on a task. This time-on-task effect was interpreted as a reduction of task engagement. To replicate it, we applied the LME model to mean theta values and mean MA9 values around its trough as a function of trial and block numbers together with two major behavioral variables, i.e., RT and congruency type. Although RTs have slightly increased only within the block (a *trial* effect), both neural measures proved affected by the time-on-task effect within each block and within the task (Table 4). Together, the MFT and MA9 presented a perfectly inverted pattern in response to these four conditions of interest (Fig. 8B). Notably, the negative relationship between these two brain markers remains even in the presence of these fixed effects in the model.

**Table 4.** Results of a follow-up LME analysis.

| Dependent | Predictor | Estim | SE | t | DF | pValue | Lower | Upper |
| --- | --- | --- | --- | --- | --- | --- | --- | --- |
| stim RT (ms) | type_incon | 89.7 | 2.0 | 44.9 | 24321 | .001 | 85.821 | 93.655 |
|  | trial | 4.5 | 0.9 | 5.1 | 24321 | .001 | 2.775 | 6.268 |
|  | block | -0.4 | 0.9 | -0.4 | 24321 | .668 | -2.108 | 1.350 |
| theta | norm RT | 0.056 | 0.007 | 8.3 | 24320 | .001 | 0.043 | 0.069 |
|  | type_incon | 0.045 | 0.015 | 3.0 | 24320 | .003 | 0.015 | 0.074 |
|  | trial | -0.032 | 0.006 | -5.0 | 24320 | .001 | -0.045 | -0.020 |
|  | block | -0.063 | 0.006 | -9.9 | 24320 | .001 | -0.076 | -0.051 |
| MA9 | norm RT | -0.065 | 0.007 | -9.8 | 24320 | .001 | -0.078 | -0.052 |
|  | type_incon | -0.034 | 0.015 | -2.3 | 24320 | .021 | -0.064 | -0.005 |
|  | trial | 0.014 | 0.006 | 2.2 | 24320 | .028 | 0.001 | 0.026 |
|  | block | 0.022 | 0.006 | 3.4 | 24320 | .001 | 0.009 | 0.034 |

Note. All fixed estimates were presented. Estim = parameter estimate. SE = standard error. DF = degrees of freedom. Lower/Upper = lower/upper confidence interval.

## 4. Discussion

Theoretical accounts indicate that theta rhythmogenesis occurs in various neural microcircuits within the medial frontal cortex (Cohen, 2014). In line with that view, previous studies conducting blind source separation analyses revealed at least two independent components of conflict-related MFT (Töllner et al., 2017; Zuure et al., 2020). Here, we aimed to investigate which independent sources contribute to the variance of the MFT amplitude. In the study, six midfrontal and two insular regions were obtained in fMRI analysis, allowing us to test for up to eight different sources of theta in two different models accounting for either conflict- or error-related variance. The results revealed only one, yet different for each analysis type, brain region showing a significant relationship with theta amplitude. The pre-response conflict-related theta was negatively correlated to the peak hemodynamic activity in MA9. In contrast, the post-response error-related theta was positively correlated to activity in aMCC. Although MFT increases in conflict and error trials are marked by disparate time-frequency characteristics (Cohen, 2014), it has been suggested that they reflect common neurophysiological mechanisms (Cavanagh et al., 2012; Cavanagh and Shackman, 2015; Nigbur et al., 2011). Our results provide strong evidence for the distinction between these two phenomena.

The positive relationship with the error-related theta and aMCC activities is in line with previous studies (Debener et al., 2005; Iannaccone et al., 2015; Nigbur et al., 2011; Wessel et al., 2012; Womelsdorf et al., 2010) and in agreement with the aMCC role in signaling surprise and the need for adaptive behavior (Ullsperger et al., 2014; Vassena et al., 2020). The lack of such a positive correlation between pre-response, i.e., conflict-related, theta and BOLD signal in preSMA or aMCC may seem surprising given the results of previous EEG source localization studies (Hanslmayr et al. 2008; Nigbur et al., 2011; Cohen and Ridderinkhof, 2013) and the finding that these brain regions show consistent activity increase with increasing response conflict. Instead, conflict-theta, as well as mean and maximum theta amplitude in epoch, showed a persistent negative relationship only to MA9. The multiple LME analysis confirmed that the deactivation in MA9 correlates with theta activity at 6 Hz around 300 ms before the response. Finally, follow-up analyses revealed a similar task-related characteristic of the two brain signals. These results provide coherent evidence that pre-response conflict theta reflects the brain activity in MA9 to a greater extent than any other brain regions in the frontal or cingulate cortex. Considering that this brain region exhibits task-related deactivation, we postulate that conflict theta reflects an inhibitory process. Although this conclusion may seem against the popular concept of theta subserving cognitive control mechanisms (Cavanagh & Frank, 2014), we further provide arguments supporting this postulate and introduce an integrative account.

The negative correlation between MFT amplitude and the BOLD signal has been reported in previous simultaneous EEG-fMRI studies. Specifically, such a relationship was found in the midline prefrontal cortex during resting state (Prestel et al., 2018; Scheeringa et al., 2008), working memory (Scheeringa et al., 2009), and decision (Algermissen et al., 2021) tasks. A study by Algermissen and colleagues (2021) was the only one to report a positive theta-BOLD correlation in the striatum. The authors argued that the striatum is too far away from the scalp and thus unlikely to be the direct neural source of MFT oscillations. Still, striato-thalamo-cortical connections may have a modulatory effect on the cortical activity that generates theta (Seifert et al., 2011; Ullsperger and Von Cramon, 2006). The question remains where the conflict, or stimulus-related, theta is generated. The theoretical account presented by Cohen (2014) suggests the medial wall of the frontal cortex for its source. Yet, the lack of positive correlation between its amplitude and other task-involved brain regions does not support them as theta generators. It is also unlikely that the source of theta, one of the most robust EEG markers, was missed in the fMRI analysis as if it did not require the hemodynamic inflow. Thus, the parsimonious explanation points to the inferences suggested by Scheeringa and colleagues (2009) that theta serves a role in task- orientedness by inhibiting irrelevant information and enabling optimal performance on a task. Our results agree with this conclusion and specify the brain region where such inhibition in the conflict task occurs, that is, the MA9.

The MA9, also referred to as the dorsal medial (or dorsomedial) prefrontal cortex, constitutes a part of the default mode network subsystem (Andrews-Hanna et al., 2014). Although its precise role is not yet clear, it seems to be linked to a wide range of self-referential functions, including mentalizing, emotional processing, and mind-wandering (Christoff et al., 2016). Consistent with this view, Durantin and colleagues (2015), using fNIRS imagining during a sustained attention task, observed a substantial activation of the medial prefrontal cortex preceding trials missed by the subjects. The authors linked this activity with mind wandering as a direct cause of omission errors. We have replicated this result using fMRI and provided further evidence that reduced task engagement is reflected by a decrease in prestimulus MFT activity. In line with that logic, suppression of mind-wandering or any other spontaneous thoughts unrelated to the task at hand during efficient cognitive control processing is highly expected. Thus, our results suggest that, like MA9, MFT is also the marker of such active inhibition.

The inhibitory nature of MFT oscillations provides a parsimonious interpretation for numerous theta-related observations. First, consistently observed and replicated in this study, the effect of positive trial-by-trial correlation between theta power and RTs (Asanowicz et al., 2021; Duprez et al., 2020) is counterintuitive the concept of cognitive control. On the one hand, cognitive control is required to select proper actions, but on the other hand, the timing of action selections is a pivotal component of its efficiency. Thus, one would expect that in the case of a homogenous type of action, the increase of cognitive control marker would speed up the action selection. Based on this assumption, Asanowicz and colleagues (2020) hypothesized an inverse relationship between the magnitude of the theta and the congruency effects and, interestingly, have found one for between-subject correlation. Second, the increase of MFT diminishes with learning (van der Cruijsen et al., 2021) and with the time on task (Arnau et al., 2021), an effect also replicated in this study. These effects can be attributed to the short-term task automatization, accompanied by segregation of the default mode network from task-related networks (Mohr et al., 2016). In other words, the task performance is less affected by spontaneous mental sketchpads as reflected by diminished theta. Finally, the great paradox of theta increasing with sleepiness and mental fatigue (Mitchell et al., 2008; Tran et al., 2020; Wascher et al., 2014) could be explained by the collective inhibition of top–down-driven metacognitive processes leading us to rest or to falling asleep.

An interesting question remains whether inhibitory processes reflected by pre-response theta activity could subserve a control mechanism. The fact that the amplitude of MFT follows the level of cognitive demands (e.g., conflict, mismatch, working memory) is not a direct proof of control mechanism. Since greater cognitive load is always accompanied by greater deactivation in midline prefrontal regions (Domagalik et al., 2012; McKiernan et al., 2003), the negative relationship between theta and activity in these regions fully accounts for theta increase. On the other hand, the phase of MFT was functionally coupled to oscillatory activity in the lateral frontal and parietal sites (Cavanagh et al., 2009b; Cohen and Ridderinkhof, 2013; Duprez et al., 2020; Hanslmayr et al., 2008). Such connectivity was interpreted as the recruitment and engagement of cognitive control mechanisms. However, the key prediction for the theta phase-dependent analysis is that certain neural computations are more likely to occur at certain theta phases (Cohen, 2014). Considering that the limited cognitive capacity creates a competition between the ongoing memory-related processes and task-related processes (Dixon et al., 2014; Lavie et al., 2004), inhibition of the former may facilitate or determine the timing of the latter. As a result, the efficacy of the neural computations in the lateral frontal and parietal cortex may be time-locked to the state of active inhibition in the MA9, that is, theta phase. Such *control by inhibition* mechanism could subserve a transition between the internally or externally focused attention as reflected by activation and deactivation of the default mode network (Buckner et al., 2008; Christoff, 2012; Clare Kelly et al., 2008). Further studies should be directed to explore and verify this account.

Finally, our findings clearly demonstrate that pre-response, conflict-related MFT, and post-response error-related MFT differ phenomenologically in frequency, latency, duration, and phase-locking to external events but also functionally in their relationship to cortical activity measured with fMRI. In contrast to conflict-related MFT, error-related MFT is clearly linked to activity increases in aMCC and the error-related negativity (Ullsperger et al., 2014). It may, thus, be questioned whether error-related MFT is indeed physiologically and functionally similar to other oscillations in the theta band. Given its short duration (roughly 1.5 wavelengths in a positive-negative-positive sequence of deflections), it might reflect transient or burst activity that happens to be in the theta range rather than an oscillation (for discussion, cf., e.g., (Luu et al., 2004b; Yeung et al., 2004). The differential links between MFT and cortical BOLD responses call for neurophysiological investigations and neuronal models shedding light on the relationship between oscillatory EEG activity and neurovascular coupling in fMRI.

## 5. Conclusions

In the study, simultaneous EEG-fMRI data were collected using two conflict tasks to couple the variance of the MFT increases related to conflict and error processing with brain activity in several frontal, insular, and cingulate subregions on a trial-by-trial level. Regarding the conflict theta, we found a consistent negative relationship with the BOLD amplitude in the MA9, a brain region showing conflict-sensitive deactivation and omission-preceding activation. Against previous claims based on EEG source estimations that the conflict theta is generated in the preSMA or aMCC, no such positive relationship was observed to either of these brain regions. However, the amplitude of post-response error-related theta was positively coupled to the activity in the aMCC in line with its established role in detecting errors and signaling the need for adaptation. MA9, on the other hand, is considered a subsystem of the default mode network that may be involved in mind-wandering. The negative relationship of this brain area with conflict theta provides a novel insight that MFT reflects an active inhibition of the self-referential processes that may distract participants from the task at hand and lead to omission error. Moreover, our results indicate that this active inhibition is attenuated with the time on task, which may serve as the neural basis for task automatization. Together, we highlight a disparate characteristic of the conflict- and error-related theta activities and emphasize that further studies investigating the cortical generators of MFT should extend the area of a search and consider the MA9 as a potential source.

## Data availability

The source data are also publicly available at https://osf.io/cx8a9/

## Code availability

All code generated for this study’s analyses are publicly available at https://github.com/ewabeldzik/thetaBOLD

## Acknowledgments

This research was supported by a National Science Centre, Poland, grant 2016/21/D/HS6/02962 (PI: Ewa Beldzik). We thank Anna Bereś, Laura Łępa, and Magdalena Wielgus for their assistance with data collection. The authors declare no conflicts of interest related to this manuscript.

## Notes

https://osf.io/cx8a9/

https://github.com/ewabeldzik/thetaBOLD

### Competing Interest Statement

The authors have declared no competing interest.

